# Competent immune responses to SARS-CoV-2 variants in older adults following mRNA vaccination

**DOI:** 10.1101/2021.07.22.453287

**Authors:** Mladen Jergović, Jennifer L. Uhrlaub, Makiko Watanabe, Christine M. Bradshaw, Lisa M. White, Bonnie J. LaFleur, Taylor Edwards, Ryan Sprissler, Michael Worobey, Deepta Bhattacharya, Janko Nikolich-Žugich

**Author notes:** To whom correspondence should be addressed - or by mail at P.O. Box 245221, Department of Immunobiology, University of Arizona College of Medicine-Tucson, 1501 N. Campbell Ave., Tucson, AZ 85724.

## Abstract

Aging is associated with a reduced magnitude of primary immune responses to vaccination and constriction of immune receptor repertoire diversity. Clinical trials demonstrated high efficacy of mRNA based SARS-CoV-2 vaccines in older adults but concerns about virus variant escape have not been well addressed. We have conducted an in-depth analysis of humoral and cellular immunity against an early-pandemic viral isolate and compared that to the P.1. (Gamma) and B.1.617.2 (Delta) variants in <50 and >55 age cohorts of mRNA vaccine recipients. We have further measured neutralizing antibody titers for B.1.617.1 (Kappa) and B.1.595; a SARS-CoV-2 isolate bearing Spike mutation E484Q. As reported, robust immunity required the second dose of vaccine. Older vaccinees manifested robust cellular immunity against early-pandemic SARS-CoV-2 and more recent variants, which remained statistically comparable to the adult group. The older cohort had lower neutralizing capacity at the first time point following the second dose, but at later time points immunity was indistinguishable between them. While the duration of these immune responses remains to be determined over longer periods of time, these results provide reasons for optimism regarding vaccine protection of older adults against SARS-CoV-2 variants and inform thinking about boost vaccination with variant vaccines.

**eTOC summary:** Vaccine responses are often diminished with aging, but we found strong responses to SARS-CoV-2 in older adults following mRNA vaccination. T cell responses were not diminished when confronted by SARS-CoV-2 variants. Neutralizing Ab were reduced but not more than those in adults.

**Graphical Abstract:** 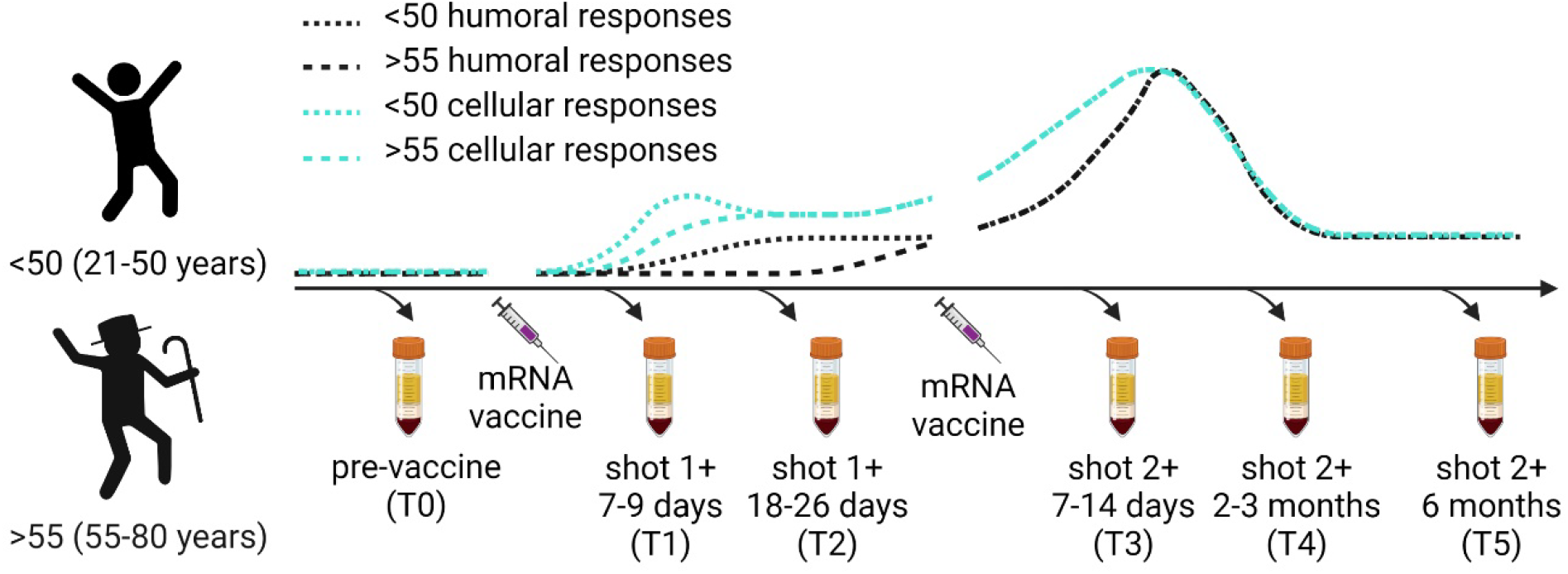

Created with BioRender.com

## Introduction

RNA viruses have a high mutation rate resulting in diverse viral populations (Robson *et al*., 2020). Since its emergence in the human population in late 2019, SARS-CoV-2 has infected more than 295 million people globally leading to the appearance of multiple new variants. Several variants of concern (VOC) quickly became dominant in their countries of identification, spreading across the globe. B.1.1.7 (Alpha) was identified in the UK in December 2020 and spread to more than 90 other countries (Karim, 2021). Other identified VOC with broad spread include B.1.351 (Beta), P.1. (Gamma) and B.1.617.2 (Delta). These lineages are transmitted more efficiently, conferring evolutionary advantage over ancestral virus (Jewell, 2021). Two mRNA vaccines based on the first published genome sequence of SARS-CoV-2 (Wuhan/Hu-1/2019), BNT162b2 (Pfizer) and mRNA-1273 (Moderna), are being deployed to reduce COVID-19 disease severity and transmission, raising the obvious question of whether they will afford similar protection against variants of concern. Reduced plasma antibody (Ab) neutralization titers against pseudovirus in mRNA vaccine recipients has been reported for multiple variants (Garcia-Beltran *et al*., 2021). Another report showed that B.1.351 (Beta) variant might be more effective in escaping humoral immunity than B.1.1.7 (Alpha) variant (Planas *et al*., 2021), as one may expect from the location of its mutations to Ab-binding regions. Despite reduced humoral immunity, T cell responses to Alpha and Beta variants were preserved in adult mRNA vaccine recipients (Geers *et al*., 2021). Similar results were obtained with adenovirus vaccine recipients, which showed 3-5 fold lower neutralizing antibody titers against Beta and Gamma variants but uncompromised T cell responses (Alter *et al*., 2021).

Efficacy and immunogenicity of many vaccines are known to be decreased in advanced age (Schmader *et al*., 2012; Young *et al*., 2017; Crooke *et al*., 2019). Fortunately, SARS-CoV-2 mRNA based vaccines were shown to be both well tolerated and highly effective in older adults (Anderson *et al*., 2020; Spector *et al*., 2020; Li *et al*., 2021). It remains unknown whether the breadth of the immune response to these mRNA vaccines will remain sufficient for protection against new viral variants. To help answer this, we have performed comprehensive analysis of antigen specific humoral and cellular immune responses in adult mRNA vaccinees of different ages. For humoral responses we compared a viral isolate representative of the ancestral virus (USA/WA1/2020, hereafter ‘WA1’) to P.1. (Gamma), B.1.617.1 (Kappa), and B.1.617.2 (Delta) and an additional local Arizona virus variant within lineage B.1.595 but, uniquely for that lineage, bearing Spike mutation E484Q (USA/AZ-WL-CVG00126, hereafter ‘AZ-E484Q’). For T cell responses, we compared WA1 with both the P.1 (Gamma) and B.1.617.2 (Delta) variants. We show that the magnitude and neutralization capacity of humoral memory to these vaccines is not reduced in our older adult cohort (>55 years) as might have been expected based on historic vaccine studies. Regarding variants, we confirm lower neutralizing antibody titers against P.1. (Gamma), B.1.617.1 (Kappa), and AZ-E484Q, but not B.1.617.2 (Delta), in our older cohort. Importantly, robust cellular immunity against WA1, P.1. (Gamma), and B.1.617.2 (Delta) was preserved in both cohorts of mRNA vaccine recipients. Overall, the effect of age on the immune response to the COVID mRNA vaccines was measurable, but minimal. It was most evident in the antibody response at the first time point after the second dose. For cellular immunity responses were essentially equivalent between the two cohorts. These results demonstrate that effective immunity in older adults is attainable with mRNA vaccines.

## Results and Discussion

A total of 40 participants were enrolled in our study before receiving either the BNT162b2 (Pfizer) or mRNA-1273 (Moderna) COVID vaccine. Blood draws were collected prior to initial dose of vaccine, 7-9, and 18-26 days after first vaccine dose, and 7-14 days, 2-3 months, and 6 months after their second dose. These time points are designated as T0, T1, T2, T3, T4, and T5 on the graphs. One participant was excluded from the study because they had a high neutralizing antibody titer and a strong antigen (Ag) specific T cell response before vaccination, indicating prior infection. Our final cohort included 23 participants under 50 years of age (<50) and 17 participants over the age of 55 (>55).

Antibody ELISA assays for the receptor-binding domain (RBD) and S2 region of Spike protein on plasma samples at each time point show similar rates and levels of seroconversion between cohorts (Figure 1A, B). Area under curve (AUC) values confirm that participants over age 55 have robust and comparable antibody responses to the younger cohort with no significant differences measured at any time point (Figure 1C). Neutralizing antibody test titers utilizing WA1 were also comparable between both cohorts across time points (Figure 2A). These data taken together are important because they demonstrate that the humoral immune response in older adults is preserved to these mRNA vaccines when tested with the virus they were generated against. We found a diminished capacity to neutralize the P.1. (Gamma), AZ-E484Q, and B.1.617.1 (Kappa) but not to B.1.617.2 (Delta) at the third time point with equivalent neutralization in memory (Figure 2A and Table 1). Our <50 cohort demonstrated reduced neutralization against AZ-E484Q, B.1.617.1 (Kappa), and B.1.617.2 (Delta) (Figure 2A and Table 1). E484Q, a mutation in the receptor binding domain (RBD) of Spike protein, is present in both B.1.351 (Beta) and P.1. (Gamma) and has already been demonstrated to impact neutralization capacity (Wang *et al*., 2021). We chose AZ-E484Q to test against vaccine induced antibodies because it has Spike E484Q and D614G mutations in common with B.1.617.1 (Kappa) but not L452R. L452R has already been shown by others to impact neutralization in pseudotyped virus systems (Li *et al*., 2020) but it remained an open question how E484Q impacts neutralization capacity (Verghese *et al*., 2021). Our use of authentic B.1.595 SARS-CoV-2, which in Spike bears only E484Q and D614G, demonstrates that E484Q also affords the virus an opportunity to escape neutralization. Understanding how this particular mutation impacts neutralization may be important during the emergence of future variants. Finally, we compared neutralizing antibody titers between vaccine brands (n=16 Moderna, n=21 Pfizer) at T3 and T4 and determined that antibody responses to the Pfizer vaccine were lower than for Moderna when tested against WA but were statistically indistinguishable when evaluated against the variant viruses (Supp. Fig.1A).

**Table 1.** Descriptive statistics of T3 for Figure 2.

|  |  | Summary | Adjusted<br>P Value |
| --- | --- | --- | --- |
| WA1/2020 | <50 vs. >55 | * | 0.0104 |
| P.1. (Gamma) | <50 vs. >55 | *** | 0.0003 |
| B.1.595 (AZ-E484Q) | <50 vs. >55 | ns | 0.9926 |
| B.1.617.1 (Kappa) | <50 vs. >55 | ns | >0.9999 |
| B.1.617.2 (Delta) | <50 vs. >55 | ns | 0.9997 |
| <50 | WA1/2020 vs. P.1. (Gamma) | ns | 0.2237 |
| <50 | WA1/2020 vs. B.1.595 (AZ-E484Q) | **** | <0.0001 |
| <50 | WA1/2020 vs. B.1.617.1 (Kappa) | **** | <0.0001 |
| <50 | WA1/2020 vs. B.1.617.2 (Delta) | **** | <0.0001 |
| >55 | WA1/2020 vs. P.1. (Gamma) | * | 0.0406 |
| >55 | WA1/2020 vs. B.1.595 (AZ-E484Q) | * | 0.0235 |
| >55 | WA1/2020 vs. B.1.617.1 (Kappa) | *** | 0.0004 |
| >55 | WA1/2020 vs. B.1.617.2 (Delta) | ns | 0.2711 |
| <50 | P.1. (Gamma) vs. B.1.595 (AZ-E484Q) | ** | 0.0044 |
| <50 | P.1. (Gamma) vs. B.1.617.1 (Kappa) | **** | <0.0001 |
| <50 | P.1. (Gamma) vs. B.1.617.2 (Delta) | *** | 0.0002 |

**Figure 1.**
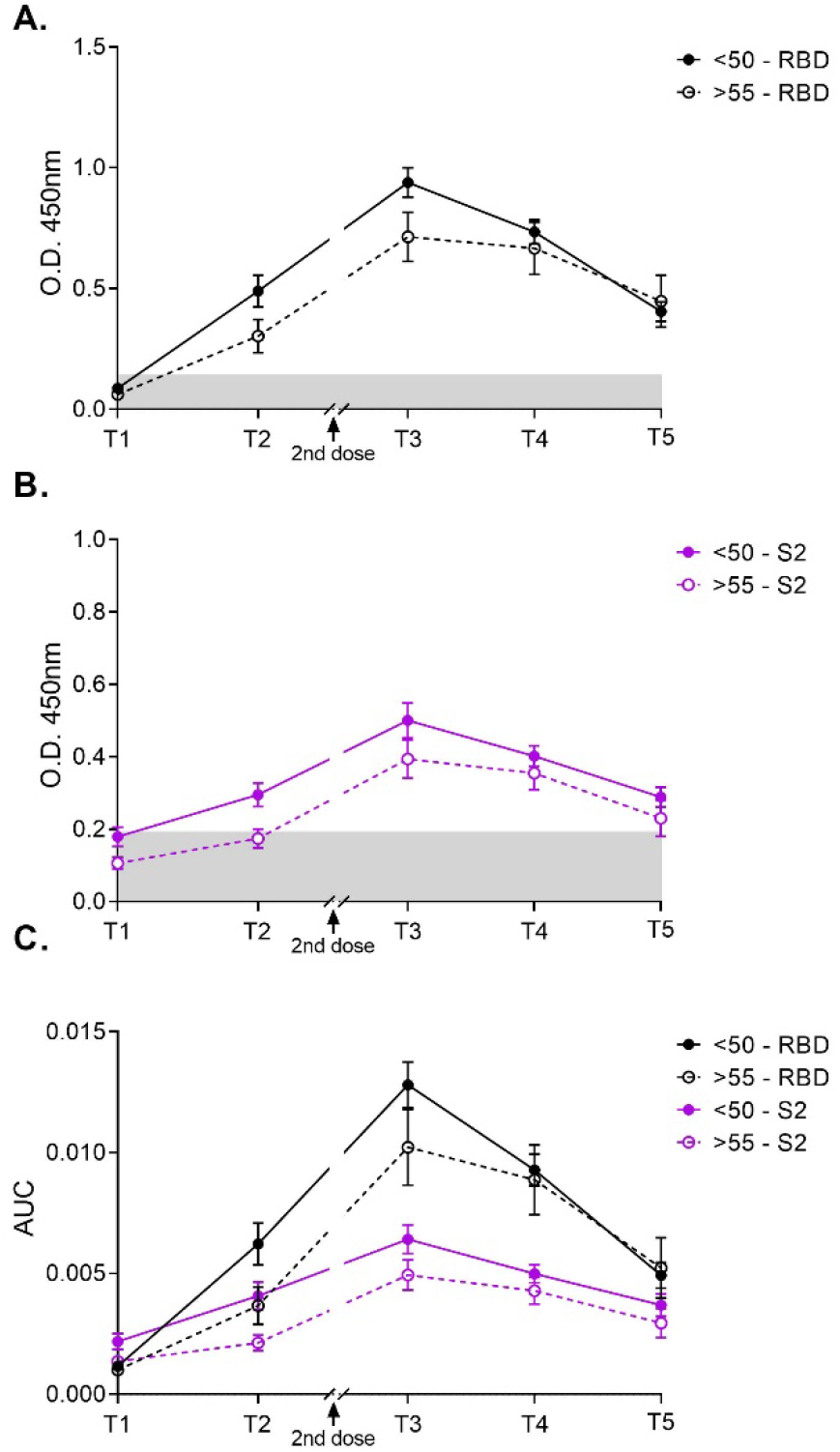
Antibody responses to mRNA vaccines are comparable regardless of age. Time points denote days 7-9 (T1) and 18-26 (T2) days post first and 7-14 days (T3), 2-3 months (T4), and 6-9 months (T5) post second vaccine dose. Semi-quantitative 1:60 serum dilution ELISA results for reactivity to RBD **(A)** and S2 **(B)** of SARS-CoV-2 Spike protein. Mean ± SEM is shown. Gray shaded area indicates positivity threshold for each assay. There is no statistical difference (Two-way ANOVA with Tukey’s multiple comparisons test) between age groups at any time point measured. **C)** Quantitative titers for RBD and S2 were calculated for each individual at each time point and are shown as area under the curve (AUC) values. There is no statistical difference (Two-way ANOVA with Tukey’s multiple comparisons test) between age groups at any time point measured.

**Figure 2.**
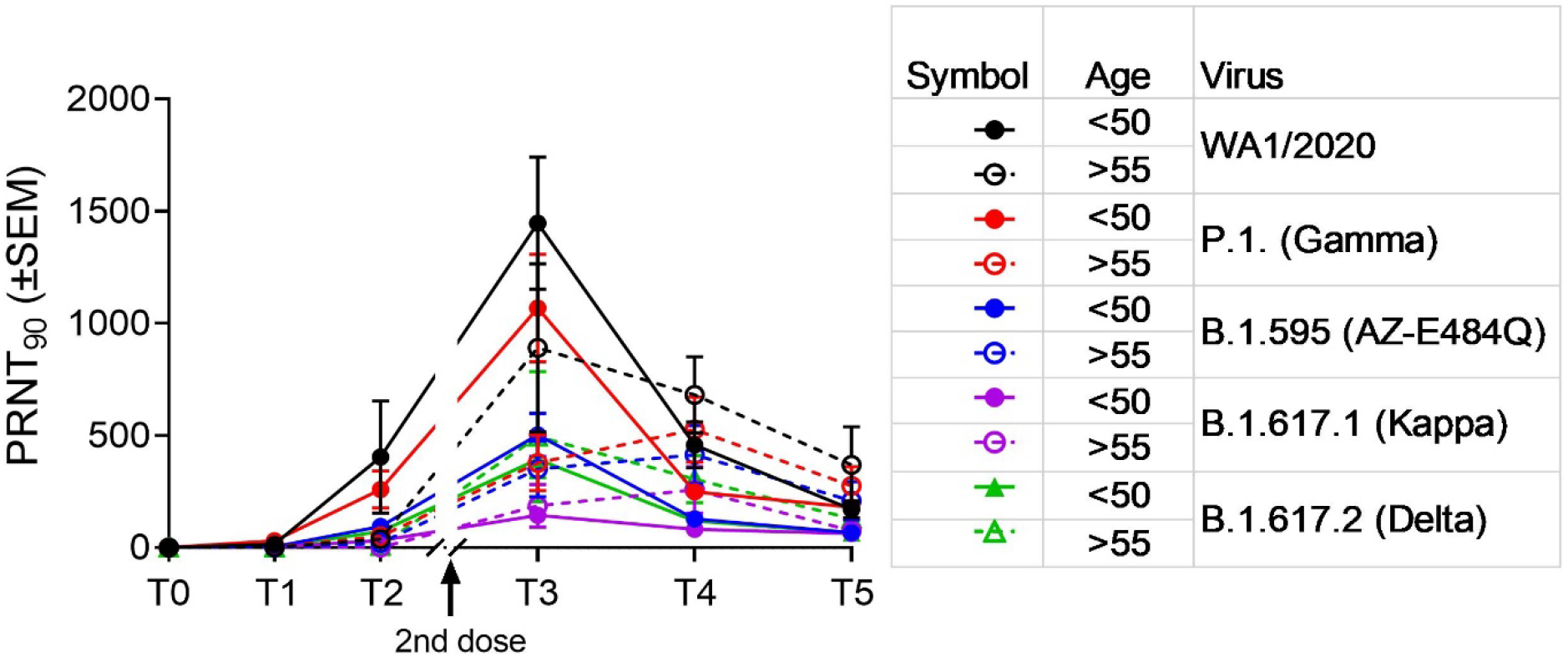
mRNA vaccines induce robust neutralization titers in adults under 50 and over 55 that is variably affected, but not abolished, by SARS-CoV-2 variants. Time points are pre-bleed (T0); days 7-9 (T1) and 18-26 (T2) days post first dose of vaccine; and 7-14 days (T3), 2-3 months (T4), and more than 6 months (T5) after second dose of vaccine. Virus neutralization assays were performed using the USA-WA1/2020, P.1. (Gamma), AZ-E484Q, B.1.617.1 (Kappa), and B.1.617.2 (Delta) isolates of SARS-CoV-2. Serial 1:3 dilutions of plasma were assayed for neutralization capacity on Vero cells. The highest dilution capable of preventing >90% of plaques was considered to be the PRNT90 value. Statistical significance was assessed by Two-way ANOVA with Tukey’s multiple comparisons test and determined significant only at time point 3 as represented in Table II.

Antibody assays are considered the gold-standard when assessing the quality of immunity after vaccination because antibodies can provide sterilizing immunity. However, the establishment of memory B cell populations is also critical to lasting vaccine efficacy as these are the resources available for response to the virus, or a variant, upon next encounter. To assess whether there is an impact of age on the formation of memory B cell populations we measured the frequency and number of circulating memory B cells specific for SARS-CoV-2 pre-vaccination and one week after booster dose by dual staining with tetramers specific for RBD and S1 using flow cytometry (Shroff *et al*., 2021) (gating strategy in Figure 3A). Both groups of participants had a small (0.1-0.3% of all B cells) but detectable population of antigen specific cells double positive for S1 and RBD tetramer (Figure 3A). This population increased as a percentage of B cells post vaccination in all but two (one from each cohort) participants with no difference between groups (Figure 3B). Similar results were obtained with S1 single-positive cells (Figure 3D) again with no difference between adult and older participants post vaccination. By multiplying percentages of tetramer positive B cells with total B cell percentage and lymphocyte counts we have calculated the absolute numbers of SARS-CoV-2 specific B cells in circulation and again observed no difference between cohorts for S1+RBD+ double positive cells (Figure 3C) or S1 single positive cells (Figure 3E). Next, we examined the phenotype of the SARS-CoV-2 specific B cells (gating strategy in Figure S1B) with respect to class switching, as it has been previously reported that aging is associated with a decline in the percentage and numbers of switched memory B cells (Frasca *et al*., 2008). To investigate this possibility, we examined differentiation and class switching of total tetramer (S1+) positive B cells by flow cytometric staining for CD27, IgM, IgD, CD21 and CD11c (representative flow cytometric gating in Figure S1A). Ag-specific B cells from adult and older participants expressed CD27 at identical levels (Figure S1C) and equal numbers of both CD27 positive and negative cells were class switched (Figure S1D,E). Adult and older participants also displayed no difference in classical memory (CD21+) phenotype among the class switched tetramer-positive cells (Figure S1F). Thus overall, we conclude that induction and differentiation of SARS-CoV-2 specific B cells through vaccination was not compromised by aging.

**Figure 3.**
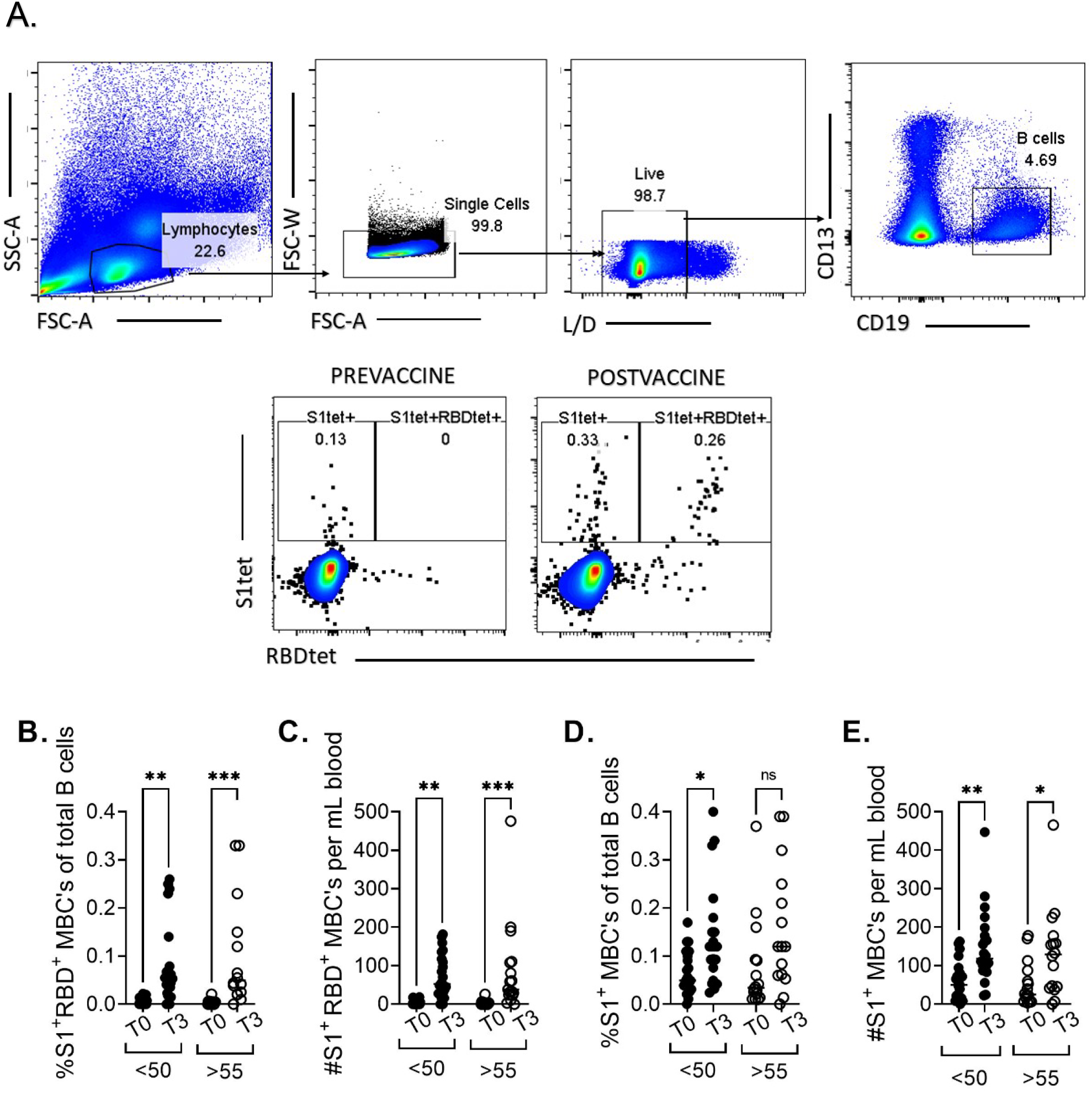

Finally, we measured antigen-specific T cells elicited by vaccinated. Given that ELISpot was previously reported to be a highly sensitive method for detection of rare antigen specific T cells (Chauvat *et al*., 2014) we simultaneously measured the number of T cells expressing costimulatory molecules CD137 and OX-40 by flow cytometry (used in several reports examining SARS-CoV-2 specific T cell immunity - (Grifoni *et al*., 2020; Rydyznski Moderbacher *et al*., 2020) (representative flow cytometry in Figure S2A) and performed IFN-γ ELISpot on PBMCs from select participants stimulated with peptide pools corresponding to the Spike protein of WA1. Confirming prior data, ELISpot proved to be a much more sensitive method for enumeration of Ag-specific T cells. We detected a statistically significant increase in ELISpots per 10^6^ PBMC’s after vaccination analyzing just 5 samples, whereas parallel flow cytometry samples showed no significant differences (Figure S2B-E). The only limitation of ELISpot, as compared to flow cytometry based enumeration, is that total T cell responses are measured without separate quantification of CD4 and CD8 responses.

We also analyzed Ag-specific T cell responses to stimulation with Spike protein peptide pools from WA1 versus two VOC by ELISpot. Participant PBMC’s were stimulated with 16-mer overlapping peptide pools corresponding to the spike protein of WA1, P.1. (Gamma) and B.1.617.2 (Delta). In accordance with previously published results (Sahin *et al*., 2020), mRNA vaccines induced a robust T cell response to WA1, the ancestral strain of SARS-CoV-2, which did not differ for Gamma and Delta variants, as evidenced by a 10-fold increase in ELISpots from post-vaccination samples stimulated by S peptide pools compared to unstimulated wells (Figure 4A). Of note, the data represented in Figure 4A shows concatenated time points for each peptide pool to demonstrate the resolution of this assay. Next, we parsed that data to compare T cell responses of both cohorts at different time points post-vaccination and subtracted the number of spots in the unstimulated wells for each sample to properly calculate the number of Ag-specific ELISpots. Data from old mice (Brien *et al*., 2009; Fulton *et al*., 2013; Jergović *et al*., 2019) showed that induction of Ag-specific T cell responses becomes delayed and decreased with age. In our data, there was a slightly lower response in the older cohort at day 7 post first dose with all three variants which was statistically significant only with the WA1 peptide pool (Figure 4B). However, after a second dose both groups had a very robust T cell response against all three viral variants examined (Figures 4C, D, E). Therefore, while we acknowledge that the primary Ag-specific T cell response is lower, and likely delayed, in some of our older participants, we conclude that the outcome after booster dose is competent T cell mediated immunity against WA1 and tested VOC in all mRNA vaccine recipients in our study. This is in agreement with our previous studies of vaccination in aged mice which showed that at least two cycles of in vivo restimulation are required for adequate ag-specific T cell response in aged animals (Uhrlaub *et al*., 2011). All of these data taken together demonstrate the rather expected blunted primary response in older adults and the improvement of this response to the levels seen in adults following a second vaccine dose. Since IFN-γ is not the only effector cytokine produced by T cells following antigen stimulation, we have additionally measured polyfunctional responses. Spike peptide pools induce a dramatic number of IFN-γ spots in comparison to unstimulated wells, but also an increase in IL-2 and GrB spots (Figure 5A). We observed no difference between the age groups in IL-2 or GrB spots (Figure 5B) in response to WA/2020 or Delta peptide pools at post-second dose time points (T3 and T4). Similarly, there was no difference between the number of polyfunctional double positive (Figure 5C) or triple positive cells (Figure 5D). We also analyzed FLUORISpot responses in recipients of mRNA vaccines from different manufacturers. We did not measure any difference in IFN-γ, IL-2 or GrB T cell responses between recipients of Moderna vs. Pfizer mRNA vaccine (Figure S3A and Figure S3B).

**Figure 4.**
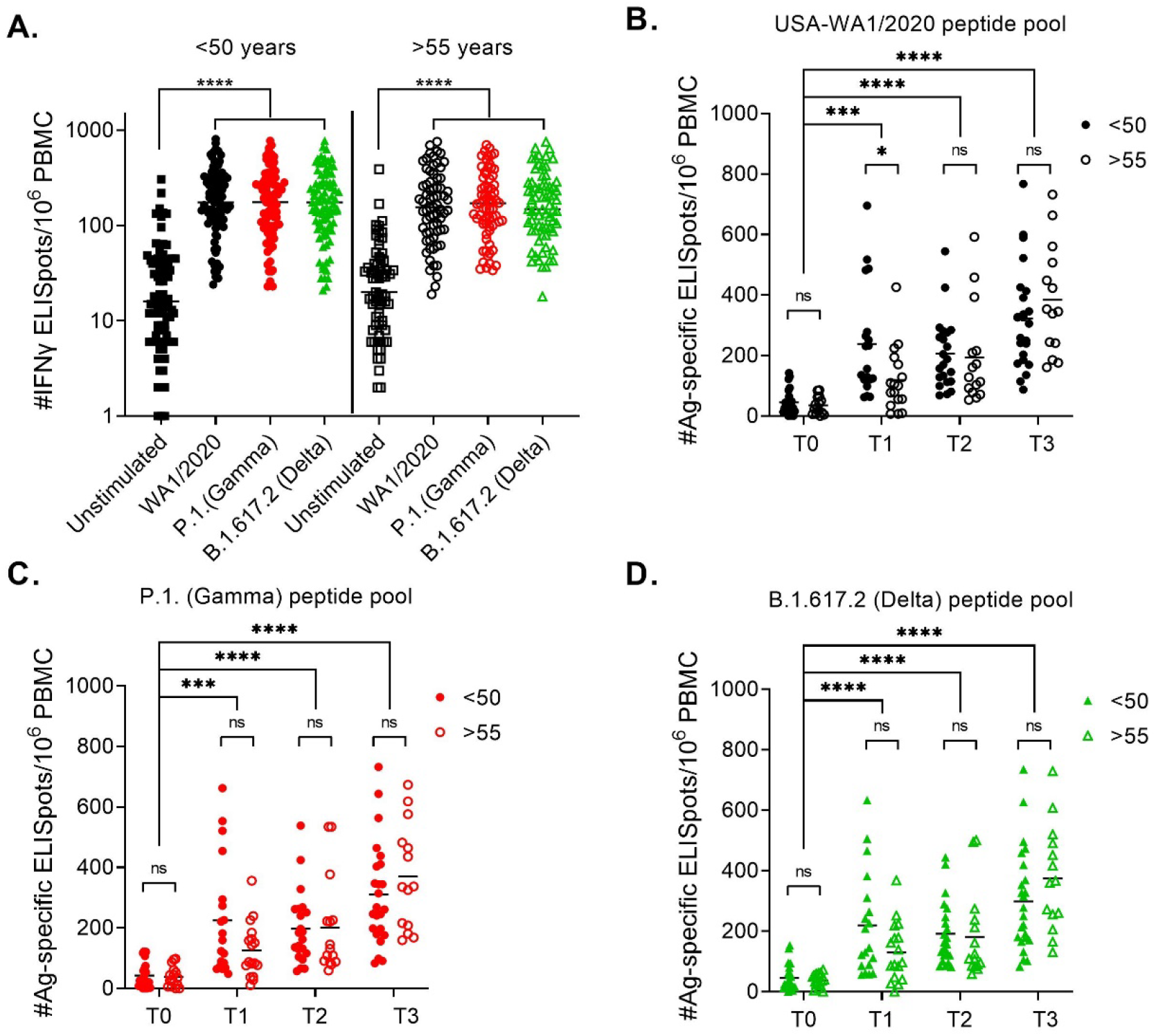
mRNA vaccines induce SARS-CoV-2 specific T cells reactive to Spike peptide pools from all three variants. **A)** 10^6^ PBMCs per well were stimulated with Spike peptide pools from USA-WA1/2020, P.1 (Gamma), and B.1.617.2 (Delta) cultured overnight in pre-coated IFN-γ ELISpot plates. All three peptide pools induced an equally strong IFN-γ response compared to unstimulated wells when all post-vaccination time points were pooled together **B)** Ag-specific ELISpot numbers were calculated by subtracting the unstimulated wells of each participant from the peptide stimulated wells. USA-WA1/2020 induced an equal number of ELISpots in <50 and >55 cohorts at later time points (T2 and T3) but the response was lower in individuals >55 yo after the first dose (T1). **C)** P.1 (Gamma) peptide pool induced a strong response at all time points post-vaccination which did not differ between <50 and >55 cohorts. **D)** Identical results were obtained with stimulation with B.1.617.2 (Delta) pool. Kruskal Wallis test with Dunn’s post hoc correction. For all statistical differences *p<0.05, **p<0.01, ***p<0.001. ****p<0.0001.

**Figure 5.**
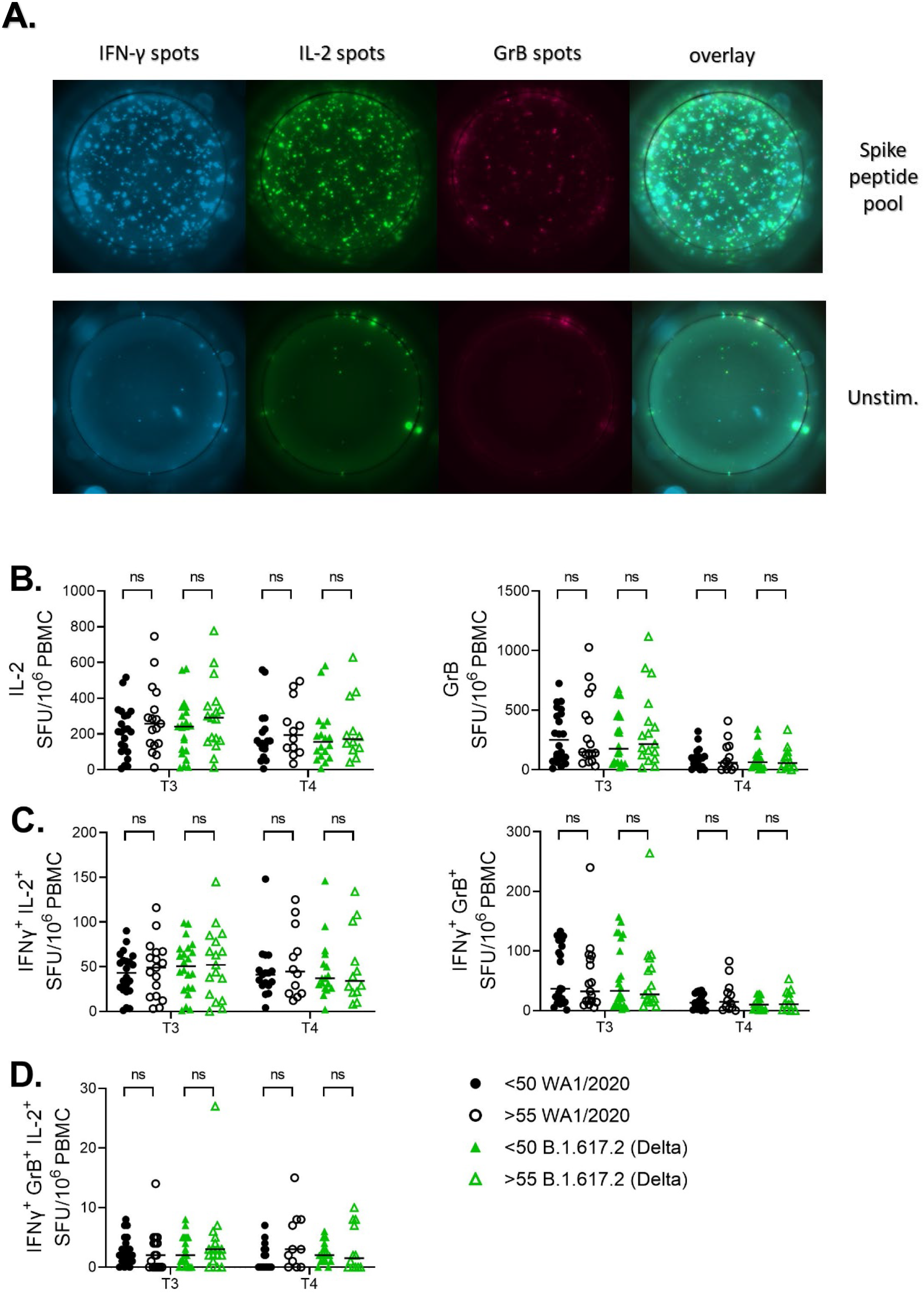
mRNA vaccines induce polyfunctional SARS-CoV-2 specific T cells in adult and older adult vaccine recipients. **A)** 10^6^ PBMCs per well were stimulated with Spike peptide pools from USA-WA1/2020 and B.1.617.2 (Delta) cultured overnight in pre-coated IFN-γ, IL-2 and GrB FLUORISpot plates. **B)** Ag-specific FLUORISpot numbers were calculated by subtracting the unstimulated wells of each participant from the peptide stimulated wells. USA-WA1/2020 and B.1.617.2 (Delta) pools induced an equal number of IL-2 and GrB spots in <50 and >55 cohorts at each time point. **C)** Similarly, numbers of double positive spots (IFN-γ+IL-2+ or IFN-γ+GrB+) did not differ between <50 and >55 cohorts with both peptide pools. **D)** Number of triple positive spots (IFN-γ+IL-2+GrB+) were also not different between the age groups and the variant peptide pools. Kruskal Wallis test with Dunn’s post hoc correction. For all statistical differences *p<0.05, **p<0.01, ***p<0.001. ****p<0.0001.

It is well established that primary immune responses wane with age and contribute to the increased susceptibility to infection experienced by older adults (reviewed in Nikolich-Žugich, 2018). There is also evidence that generation of immune memory in older adults is reduced, but not maintenance of memory. Studies with multiple pathogens have shown decreased T cell receptor repertoire with age (Yager *et al*., 2008; Smithey *et al*., 2018) which could mean easier escape from existing immunity for pathogen variants. All of the above-mentioned findings warrant an extensive and long-term monitoring of immunity in SARS-CoV-2 vaccinees, especially those over age 55.

The emergence of SARS-CoV-2 impacted older adults especially hard with more than 80% of deaths in those over age 65 (Lithander *et al*., 2020) and mortality rates rising sharply above the age of 55 (Yanez *et al*., 2020). Vaccination reduces these rates dramatically (Haas *et al*., 2021) demonstrating that the principles of immunology hold true for this virus and that immune memory is what our species is lacking. Well tolerated and effective, SARS-CoV-2 mRNA vaccines induce potent humoral and cellular immune responses (Anderson *et al*., 2020; Sahin *et al*., 2020). Their deployment also offers an opportunity to establish, in a truly immune naive population, correlates, and contours, of protective immunity. Recent elegant studies in rhesus macaques have shown that even sub-sterilizing neutralizing antibody titers are protective and decrease SARS-CoV-2 severity (McMahan *et al*., 2021). The same research study demonstrated a protective role for T cell memory responses and this is supported by human research showing that accumulation of oligoclonal CD8 T cells in bronchoalveolar lavage fluid inversely correlated with disease severity (Liao *et al*., 2020). Recently, Collier et al. analyzed 102 partially and 38 fully vaccinated subjects and concluded that subjects >80 years of age produced lower primary, but not secondary antibody neutralizing responses, including those against variants (Collier *et al*., 2021). They did not analyze T cell immunity against variants and, somewhat curiously, did not observe increased T cell responses following the second dose in their older groups (>80). These authors argue that older adults – those over 80 – remain vulnerable at least until they receive the second vaccine dose. Our results agree with these conclusions with regard to antibody immunity, and suggest that T cell immunity in response to mRNA vaccines is robust in older adults and against variants, even though we did not analyze subjects in the octogenarian bracket. Clinical efficacy of the mRNA vaccines in protecting older adults has been strong, consistent with both our data and those by Collier et al (Collier *et al*., 2021). The decrease in antibody titer when challenged with more recent SARS-CoV-2 variants does suggest that the breadth of immunity may be narrower in advanced age; a challenge that can be met with booster doses of heterologous sequence virus. Further studies on the durability and breadth of protection by current and future heterologous vaccines in older adults will be necessary to answer these and other germane questions on their immunity and SARS-CoV-2 protection in older adults.

## Materials and methods

### Study participants

This study was approved by the University of Arizona IRB (Protocol#2102460536). We collected blood from 23 adults <50 years old and 17 above 55 years old. Demographics are provided in Table II. Samples for all participants were collected prior to vaccination, one week after first dose (mRNA vaccine: Pfizer N=23; Moderna N=17), day before booster dose and 7-10 days after booster. Many participants were also sampled at 3 or 6-9 months following first dose. The time points are labeled T0, T1, T2, T3, T4, and T5 in all graphs. Blood for complete blood count was collected in BD vacutainer with EDTA and submitted to Sonora Quest Laboratories (Arizona). Blood for peripheral blood mononuclear cells (PBMCs) and plasma was collected in BD Vacutainer with sodium heparin. Plasma was separated by centrifugation at 1100 rpm for 10 minutes and PBMC was isolated from the buffy coat by Ficoll-Paque PLUS (GE Healthcare) and cryopreserved in fetal calf serum + 10% DMSO.

**Table II.**
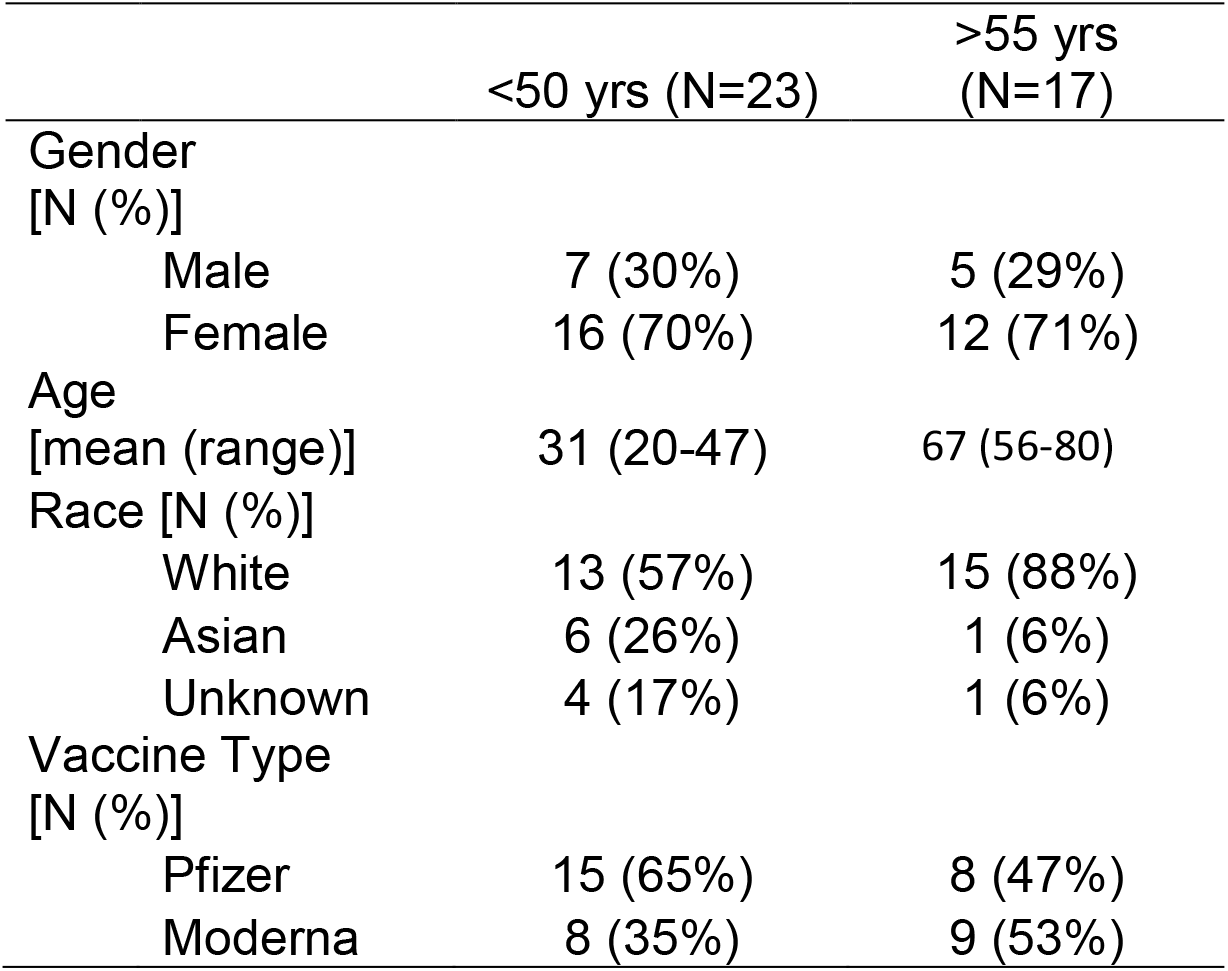
Demographic characteristics of the cohort.

### Virus

SARS-Related Coronavirus 2 (SARS-CoV-2), Isolate USA-WA1/2020, was deposited by Dr. Natalie J. Thornburg at the Centers for Disease Control and Prevention and obtained from the World Reference Center for Emerging Viruses and Arboviruses (WRCEVA). P.1. (Gamma) variant was obtained through BEI Resources, NIAID, NIH: SARS-Related Coronavirus 2, Isolate hCoV-19/Japan/TY7-503/2021, NR-54982, contributed by National Institute of Infectious Diseases. B.1.617.1 was received from BEI (NR-55486 SARS-CoV-2, Isolate hCoV-19/USA/CA-Stanford-15_S02/2021 (Kappa Variant) B.1.617.1 G EPI_ISL_1675223). B.1.617.2 was received from WRCEVA. Documentation says it was isolated from a patient at a Galveston County (UTMB) hospital; collected 7/6/2021 and passaged once on Vero cells. Strain designation GNL-1205. It has silent mutation C24208T by Sanger sequencing. AZ-E484Q, a B.1.595 virus (GISAID: EPI_ISL_765942) was grown from a nasopharyngeal swab on Calu-3 cells for 48 hours. AZ-E484Q has Spike D614G and E484Q mutations but has not emerged as a variant of concern. Stocks of SARS-CoV-2 were generated as a single passage from received stock vial (USA-WA1/2020, P.1., B.1.617.1, B.1.617.2) or primary culture (B.1.595) on mycoplasma negative Vero cells (ATCC #CCL-81) at a MOI of 0.005 for 48 h. Supernatant and cell lysate were combined, subjected to a single freeze-thaw, and then centrifuged at 3000RPM for 10 min to remove cell debris.

### Plaque Reduction Neutralization Test

A plaque reduction neutralization test (PRNT) for WA1 and SARS-CoV-2 variants was performed as described in Ripperger *et al*., 2020. Briefly, mycoplasma negative Vero cells (ATCC # CCL-81) were plated in flat bottom, 96 well, tissue culture plates and grown overnight. Three-fold serial dilutions (1:20 - 1:43,740) of plasma/serum samples were incubated with 100 plaque forming units of virus for 1 h at 37°C. Plasma/serum dilutions plus virus were transferred to the cell plates and incubated for 2 h at 37°C, 5% CO2 then overlayed with 1% methylcellulose in media. After 96 hours, plates were fixed with 10% Neutral Buffered Formalin for 30 min and stained with 1% crystal violet. Plaques were imaged using an ImmunoSpot Versa (Cellular Technology Limited) plate reader. The serum/plasma dilution that contained 10 or less plaques was designated as the NT90 titer.

### ELISpot assays

T cell specific immunity to peptide pools corresponding to Spike, Nucleocapsid, and Matrix proteins were measured as previously described (Shroff *et al*., 2021). Briefly, frozen PBMCs were thawed and rested overnight incubated in 24-well plates overnight at 37°C with 5% CO2. Following day cells were stimulated in X-VIVO 15 media with 5% male human AB serum containing ∼1 nmol of peptide pool corresponding of SARS-CoV-2 Prot S of US-WA (wild-type), Gamma (P.1./B1.1.28) and Delta (B.1.617.2) variants or positive control anti-CD3 mAb CD3-2 from or blank media as negative control. Overlapping 16mer peptide pools were purchased from 21st century Biochemicals,Inc. Cell suspensions were transferred to pre-coated Human IFN-γ ELISpot plus kit (Mabtech) and developed after 20h according to manufacturer instructions. Spots were imaged and counted using an ImmunoSpot Versa (Cellular Technology Limited).

### Antibody ELISA

Serological assays were performed exactly as previously described in Shroff *et al*., 2021). Area under the curve (AUC) values were calculated using R version 4.0.3 (R Core Team, 2020) and the DescTools package (Signorelli, 2021).

### Flow cytometry

Cryopreserved PBMC (2-5 × 10^6^/sample) were thawed in prewarmed RPMI-1640 with L-glutamine (Lonza) + 10% FCS. Thawed PBMCS were rested overnight at 37 ^°^C in X-VIVO 15 Serum-free Hematopoietic Cell Medium (Lonza) supplemented with 5% human Ab serum. Cells were stained with surface antibodies in PBS (Lonza) + 2% FCS, and then stained with the live dead fixable blue dye (Thermofisher). B cell tetramers were assembled by mixing 100 μg ml−1 of C-terminal AviTagged RBD or S1 (ACROBiosystems) with 100 μg ml−1 of streptavidin-PE (eBiosciences) or streptavidin-BV421 (BioLegend), respectively, at a 5:1 molar ratio in which 1/10 of the final volume of streptavidin was added every 5 min. Samples were stained for 1h at 40 C. Samples were acquired using a Cytek Aurora cytometer (Cytek) and analyzed by FlowJo software (Tree Star).

### FLUORISpot assays

Cryopreserved PBMC (5 × 10^6^/sample) were thawed in prewarmed RPMI-1640 with L-glutamine (Lonza, Basel, Switzerland) + 10% FCS. Thawed PBMCS were rested for 3-4h at 37°C in X-VIVO 15 Serum-free Hematopoietic Cell Medium (Lonza) supplemented with 5% human Ab serum. Cells were then stimulated with ∼1 nmol of peptide pool corresponding to SARS-CoV-2 spike (S) of US-WA (ancestral), P.1. (Gamma), or B.1.617.2 (Delta). Peptides were 16mer peptide pools, overlapping by 10 amino acids, purchased from 21st century Biochemicals Inc. Cell suspensions were transferred to pre-coated Human IFN-γ, IL-2, Granzyme-B (Gz-B) FLUORISpot kit plates (Mabtech, Inc.) and developed after 48h according to manufacturer instructions. Spots were imaged and counted using a Mabtech Iris Fluorispot reader (Mabtech).

### Statistical analysis

SPSS and Graph Pad Prism were used for statistical analysis. Upon inspection of data distribution by Shapiro-Wilks normality test group differences were calculated by Mann Whitney U-test or Kruskal-Wallis test with Dunn’s post-hoc correction. For comparisons where groups means were compared over differing time points two-way ANOVA with Tukey post hoc correction for multiple comparisons. For all statistical differences *p<0.05, **p<0.01, ***p<0.001. ****p<0.0001.

## Data Availability

All data is available upon request.

---

**Figure S1.**
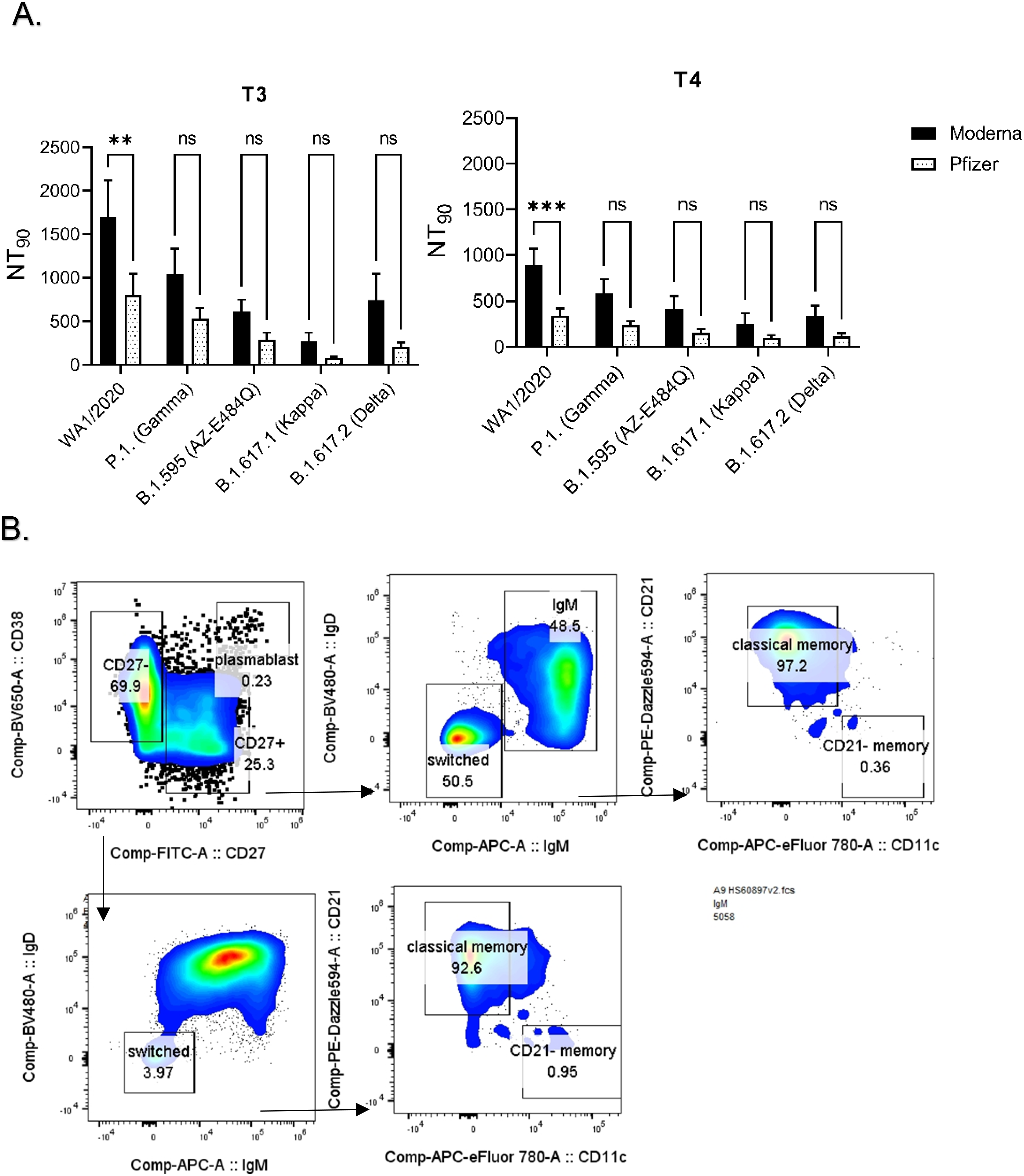
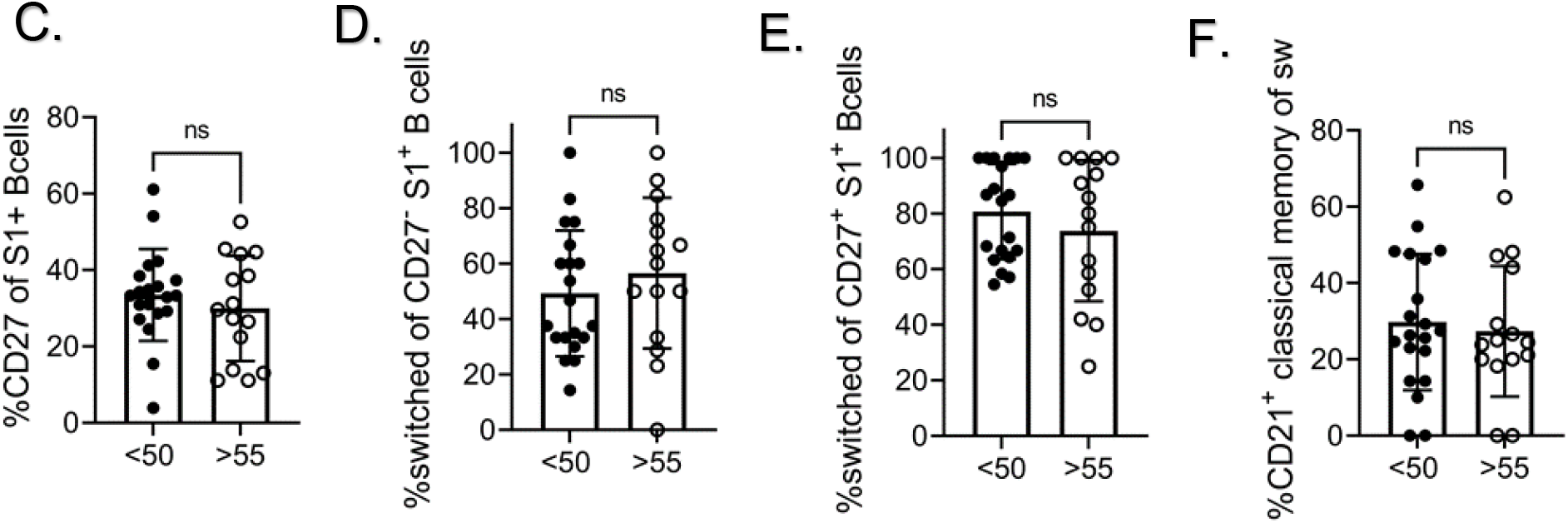
SARS-CoV-2 specific memory B cells differentiate equally in <50 and >55 cohorts. **A)** Lower neutralization titers in recipients of Pfizer’s mRNA vaccine. **B)** Representative flow cytometric gating to examine B cell differentiation. **C)** Percentage of CD27 expressing B cells among tetramer positive cells was equal between cohorts. **D & E)** Equal percentage of SARS-CoV-2 specific B cells were class switched among CD27 positive and negative cells between <50 and >55 cohorts. **F)** Expression of CD21 (Classical memory) was also equal between <50 and >55 cohorts. Mann-Whitney U test, for all statistical differences *p<0.05, **p<0.01, ***p<0.001. ****p<0.0001.

**Figure S2.**
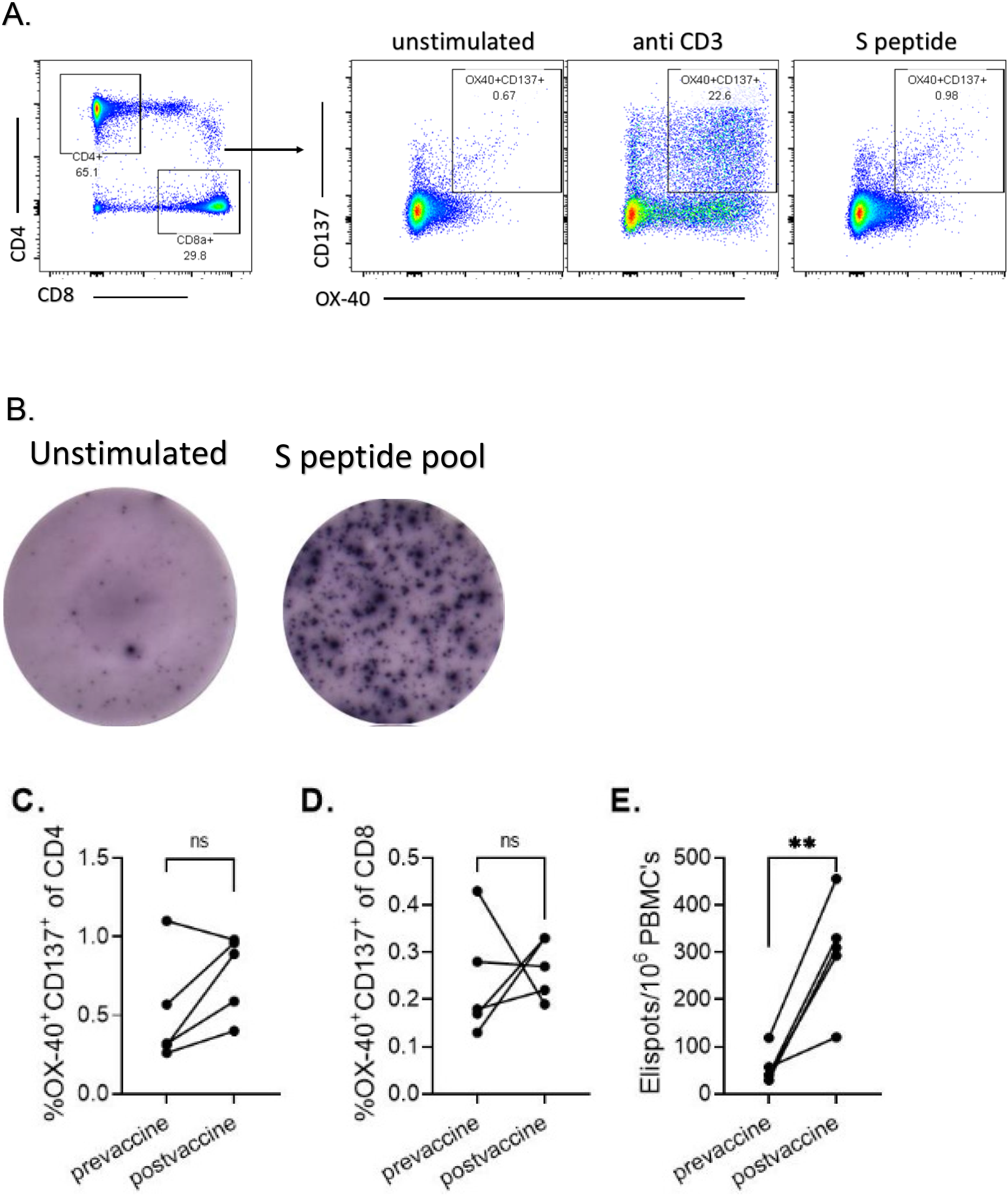
ELISpot is more specific method for enumeration of SARS-CoV-2 specific T cells. **A)** Representative flow cytometric gating for identifying CD137+OX-40+ double positive CD4 and CD8 T cells. **B)** Representative unstimulated and Spike protein peptide pool stimulated ELISpot wells. PBMCS from 5 participants were stimulated overnight with USA-WA1/2020 spike peptide pool. SARS-CoV-2 specific T cells were enumerated by either flow cytometry **(C**,**D)** or ELISpot **(E)**. ELISpot proved to be a more sensitive method with statistically significant difference before and after vaccination with N=5. Mann-Whitney U test, for all statistical differences *p<0.05, **p<0.01, ***p<0.001. ****p<0.0001.

**Figure S3.**
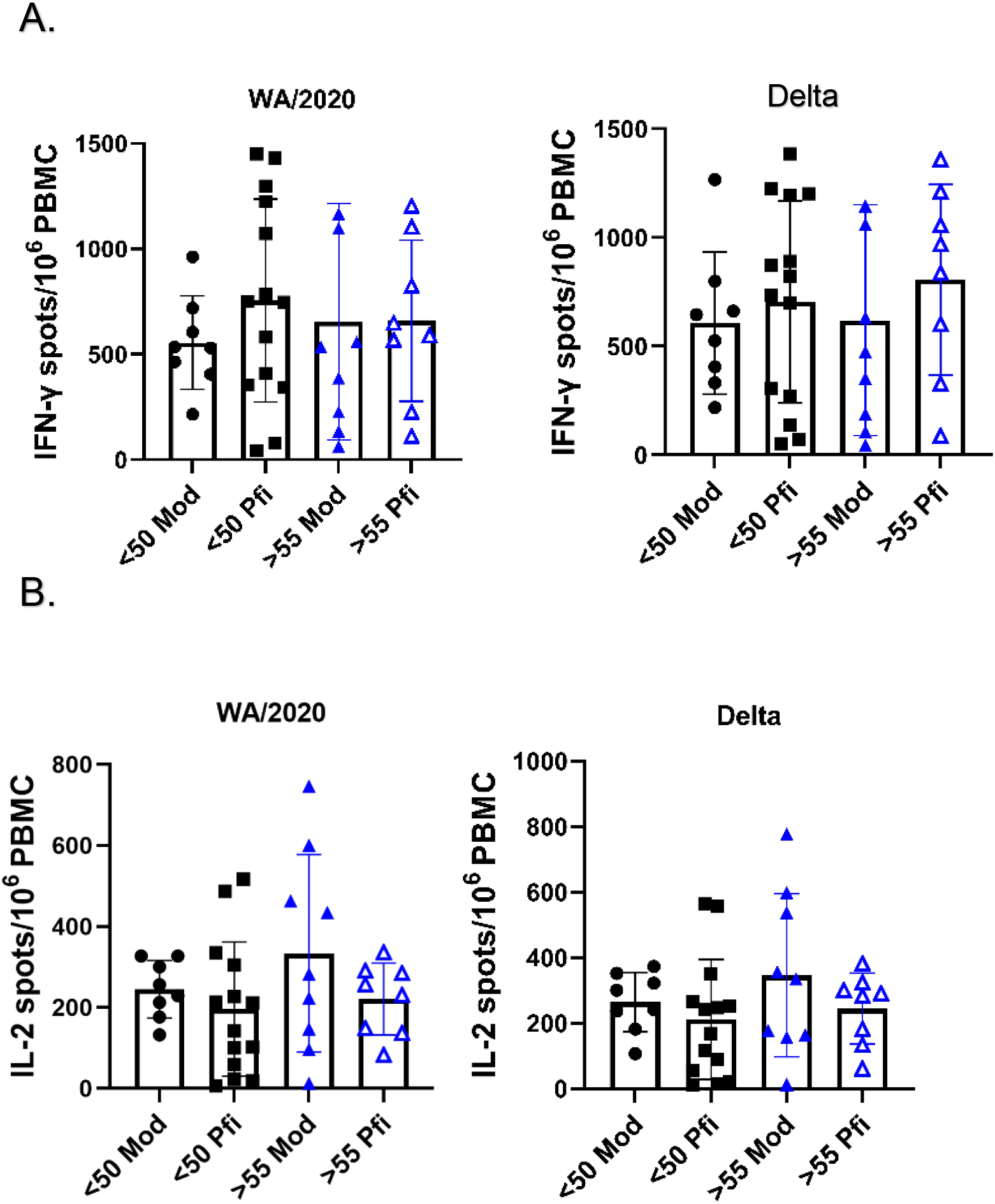
No difference in T cell responses between recipients of different mRNA vaccine brands. **A)** Number of IFN-γ spots was the same between recipients of Moderna and Pfizer vaccines at T4 time point **B)** Number of IL-2 spots was the same between recipients of Moderna and Pfizer vaccines at T4 time point for. Kruskal-Wallis test with Dunn’s post-hoc correction. For all statistical differences *p<0.05, **p<0.01, ***p<0.001. ****p<0.0001.

